# SleepSatelightFTC: A Lightweight and Interpretable Deep Learning Model for Single-Channel EEG-Based Sleep Stage Classification

**DOI:** 10.1101/2024.08.02.606301

**Authors:** Aozora Ito, Toshihisa Tanaka

**Author notes:** Corresponding author: Toshihisa Tanaka.

## Abstract

Sleep scoring by experts is necessary for diagnosing sleep disorders. To this end, electroencephalography (EEG) is an essential physiological examination. As manual sleep scoring based on EEG signals is time-consuming and labor-intensive, an automated method is highly desired. One promising automation technology is deep learning, which has performed well or better than experts in sleep scoring. However, deep learning lacks adequate interpretability, which is crucial for ensuring safety and accountability, especially for complex inference processes. We propose SleepSatelightFTC, a lightweight model that achieves comparable performance to state-of-the-art models with only one-third of their parameters. Based on the rules for sleep scoring, self-attention is applied to each of the time- and frequency-domain inputs, a raw EEG signal and its amplitude spectrum. The simple method of continuously connecting the intermediate outputs of the epoch-wise model has resulted in a highly lightweight architecture. On the Sleep-EDF-78 dataset, our model achieves an accuracy of 84.8% and a kappa coefficient of 0.787 while requiring significantly fewer parameters (0.47*×* 10^6^) compared to existing models (1.3–4.54*×* 10^6^). The visualization of feature importance obtained from self-attention confirms that the proposed model learns representative waveform features, including K-complexes and sleep spindles.

## I. INTRODUCTION

SLEEP is critical to physical and mental well-being. Long-term sleep disruption is associated with an increased risk of a variety of diseases, including cardiovascular disease, obesity, and diabetes mellitus [1], [2]. Even acute sleep deprivation adversely affects daily life, including impaired judgment and cognitive abilities [2].

Polysomnography (PSG) is needed for diagnosing sleep disorders. PSG involves simultaneous recordings of various biological signals throughout the night, including electroencephalograpy (EEG), electrooculography, and electromyography signals. Based on recorded signals, experts can evaluate the sleep stages in 30 s epochs according to the American Academy of Sleep Medicine (AASM) sleep scoring manual [3]. AASM defines five sleep stages: Wake (W), rapid eye movement (REM or R), and three non-REM stages (N1, N2, N3) [3]. A complete sleep cycle takes roughly 90 to 110 minutes, with each cycle typically comprising five stages: W (∼5% of sleep), N1 (∼5%), N2 (∼45%), N3 (∼25%), and REM (∼25%) [4]. Wake is characterized by high-frequency beta waves, while N1, a transitional stage, is marked by low-voltage theta waves. N2, the most prevalent stage, features sleep spindles and K-complexes, which are crucial for sensory processing. N3, deep sleep, is dominated by high-amplitude delta waves, essential for physiological restoration. REM sleep exhibits low-amplitude beta waves, similar to wakefulness, but includes rapid eye movements and muscle atonia [4]. The decision rules to identify the sleep stages are based on features specific to each stage in the acquired biological signals. In particular, temporal and frequency features of EEG signals are listed in the scoring manual as features that define each stage in all five sleep stages. The recommended EEG electrode positions for PSG are frontal, central, and occipital. In addition, some sleep stages can be determined by considering context from adjacent stages.

Manual sleep scoring is time-consuming and labor-intensive for knowledgeable and experienced experts [5]. Even a skilled expert can take up to 2 hours to score approximately 8 hours of sleep data [6]. Manual sleep scoring impedes suitably handling the millions of patients with sleep disorders [7], rendering automated scoring required. Automation of sleep scoring is expected to reduce the burden on specialists by several thousand hours per year [8].

Sleep scoring based on EEG follows predefined rules, making it suitable for automation using machine learning [8]. Machine learning models have been proposed to determine the sleep stage from physiological signals such as EEG, electrooculography, and electromyography [9]–[11]. Additionally, several models have been proposed to determine sleep stages based on only a single-channel EEG. Automatic sleep staging based on single-channel EEG can simplify PSG, thereby reducing the burden on physicians and patients. For these models, various types of inputs have been utilized, including raw EEG signals [12], [13] and spectrograms [14]– [16]. Deep learning models for sleep staging based on single-channel EEG have achieved equivalent or better performance than expert judgment [17].

However, such deep learning models lack interpretability due to the complexity of inference compared with classical machine learning models, hindering experts to judge the validity of inferences. Classical machine learning methods for sleep stage classification are relatively easy to interpret in terms of feature and sample contributions, but their classification performance is generally inferior to deep learning methods [18]. This lack of interpretability may increase the difficulty to identify reasons underlying misclassifications, the inability to explain the model decisions to patients and healthcare providers, and potential bias in model predictions. The interpretability of models is also essential to ensure safety, ethics, and accountability [19]. The interpretation of incorrect inferences can also contribute to improve the model performance.

The self-attention mechanism [20] allows to interpret machine learning models through the visualization of feature importance. Self-attention automatically adjusts the feature importance in learning, indicating strong attention to specific data features. SleepTransformer [15] applies self-attention to spectrograms, while cross-modal transformers apply attention to EEG and EOG signals separately, enabling the interpretation of inference based on their weights. In this paper, we extend this idea by applying self-attention to the time- and frequency-domains of EEG. This allows us to quantify the contribution of each time point and frequency to the inference. We propose a single-channel EEG-based sleep stage classification model called SleepSatelightFTC that includes self-attention for interpretability. Based on the rules for sleep scoring, self-attention is applied to each of the time- and frequency-domain inputs, a raw EEG signal and its amplitude spectrum. The amplitude spectrum has lower dimensionality compared to spectrograms and can concisely represent the overall frequency characteristics of the signal. This is expected to improve interpretability while reducing the dimensionality of the input data to the model, enabling the development of lightweight models through a reduction in the number of parameters and computational cost. To reflect the sleep context, we apply transfer learning to continuous epoch data.

## II. METHODS

### A. DATASET AND PREPROCESSING

We used a public dataset, Sleep-EDF Database Expanded [21], [22] in this study. Sleep-EDF Database Expanded has two versions (2013 and 2018) and two subsets (Sleep Telemetry and Sleep Cassette). Most research on sleep stage classification has used either the 2013 or 2018 version of Sleep Cassette. We used both versions separately. The 2013 Sleep Cassette (Sleep-EDF-20) consists of PSG data over 39 nights acquired from 20 healthy participants (10 males and 10 females) aged 25–34 years. The 2018 Sleep Cassette (Sleep-EDF-78) consists of PSG data over 153 nights acquired from 78 healthy participants (37 males and 41 females) aged 25– 101 years. The PSG data were annotated by an expert according to the R & K manual [23], which defines six sleep stages. Only the Fpz–Cz EEG channel was used for evaluation. The sampling frequency of the acquired EEG signals was 100 Hz. The preprocessing pipeline followed standard procedures for EEG-based sleep stage classification. First, an anti-aliasing brick-wall filter was applied with a Nyquist frequency of 25 Hz before downsampling to 50 Hz. No additional bandpass filtering or denoising was performed. To ensure consistency in data labeling, we excluded epochs marked as ‘‘MOVEMENT’’ or ‘‘?’’, which indicate excessive body motion artifacts or noise, as per the R & K manual used in the Sleep-EDF dataset. These labels denote epochs where sleep staging was deemed unreliable by expert annotators. This exclusion approach aligns with previous studies, including TinySleepNet [17]. Unlike some previous works, no additional normalization or rescaling procedures (e.g., z-score normalization) were applied to the EEG signals. To adhere to the manual of the American Academy of Sleep Medicine, which defines five sleep stages, all the N4 labels were merged into the N3 label. Finally, we extracted the section from 30 min before the start of sleep to 30 min after the end of sleep to exclude periods unrelated to sleep. The number of epochs per label in Sleep-EDF-20 and Sleep-EDF-78 are listed in Table 1.

**TABLE 1:**
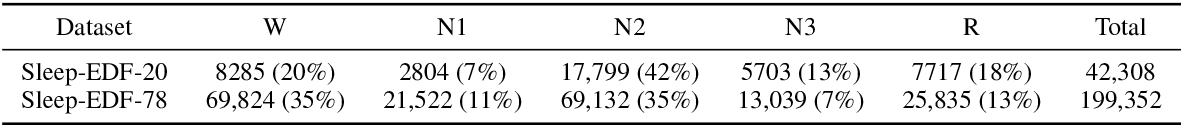
Number of epochs per label in Sleep-EDF-20 and Sleep-EDF-78. Numbers in parentheses indicate the percentage of each label in the total data set.

| Dataset | W | N1 | N2 | N3 | R | Total |
| --- | --- | --- | --- | --- | --- | --- |
| Sleep-EDF-20 | 8285 (20%) | 2804 (7%) | 17,799 (42%) | 5703 (13%) | 7717 (18%) | 42,308 |
| Sleep-EDF-78 | 69,824 (35%) | 21,522 (11%) | 69,132 (35%) | 13,039 (7%) | 25,835 (13%) | 199,352 |

### B. PROPOSED SLEEPSATELIGHTFTC MODEL

Most rules for sleep scoring are based on temporal or frequency features extracted from EEG signals. Accordingly, we propose a sleep stage classification model that processes EEG epochs as shown in Fig. 1. High-frequency activity (gamma TABLE 1: Number of epochs per label in Sleep-EDF-20 and Sleep-EDF-78. Numbers in parentheses indicate the percentage of each label in the total data set. waves with frequency *>* 30 Hz) in an EEG signal is likely related to the sleep-wake cycle but represents less than 1% of the total power spectrum [24]. Furthermore, gamma waves are likely to be affected by artifacts and noise. Thus, we down-sample the EEG signals to 50 Hz to establish a time-domain input. By reducing the input size of the model, we expect to reduce the number of parameters in the overall model. In spectral analysis for identifying differences in sleep EEG signals, the multi-taper method outperforms the single-taper method [25]. Thus, we calculate the amplitude spectrum in 0–25 Hz by applying the multi-taper method [26] to the EEG signals. The amplitude spectrum is expressed in decibel– microvolts (dB µV) by taking the logarithm to establish the input in the frequency domain. The proposed model applies\ self-attention to each input. Self-attention highlights the input that contributes to inference and allows to visualize the attention strength. Thus, self-attention is expected to enable the visualization of the model features in the time and frequency domains during training.

**FIGURE 1:**
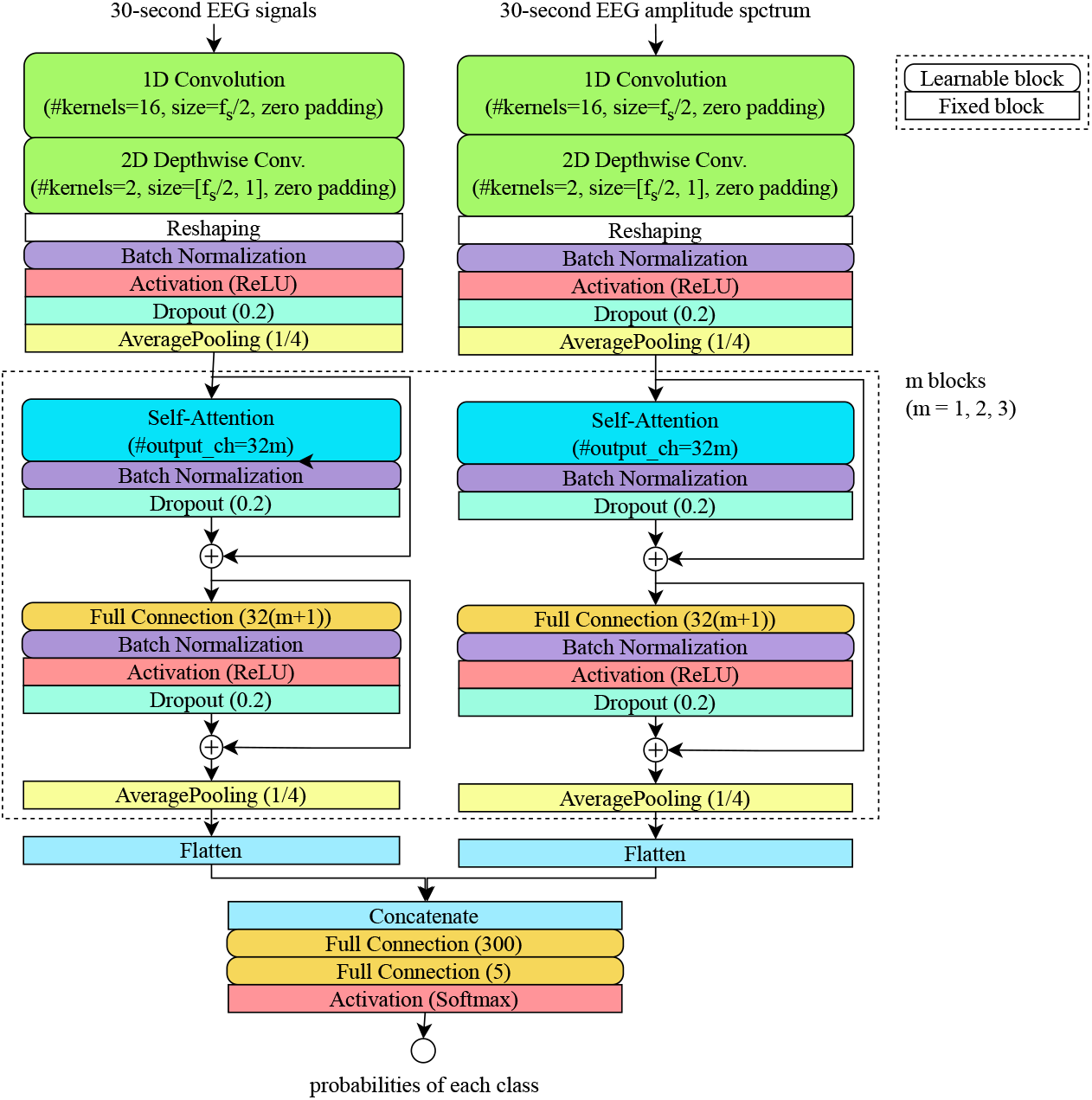
Epoch-wise classification model. The time-domain input is a 30 s raw EEG signal with 1500 samples (sampling rate *f*_*s*_ of 50 Hz), and the frequency-domain input is the 0–25 Hz amplitude spectrum with 750 samples. The model consists of convolutional layers with 16 kernels of size *f*_*s*_*/*2 for both the time- and frequency-domain inputs, followed by 2D depthwise convolutional layers with 2 kernels per input matrix, batch normalization, rectified linear unit (ReLU) activation, dropout of 0.2, and average pooling of 1/4. The model then applies *m* blocks (*m* = 1, 2, 3) of simple self-attention layers, dense layers, batch normalization, activation, dropout of 0.2, and average pooling of 1/4. The self-attention layer has 32*m* output channels. The outputs of the two input branches are flattened, concatenated, and passed through two fully connected layers with 300 and 5 units, followed by softmax activation.

Sleep shows long-term context, and most existing models for sleep stage classification employ architectures that consider the context before and after every evaluated epoch, such as RNNs [14], [27] and Transformers [15], [28]. The proposed sleep stage classification model, SleepSatelightFTC, has the architecture shown in Fig. 2. This model applies transfer learning to the base epoch-wise classification model shown in Fig. 1. The outputs of fully connected layers in the epoch-wise classification model are combined to obtain sequential epochs and used as input for transfer learning. The model output is a one-epoch sleep stage, and its input is an odd number of sequential epochs centered on the target epoch for inference. The number of epochs is selected as an odd number between 3 and 29.

**FIGURE 2:**
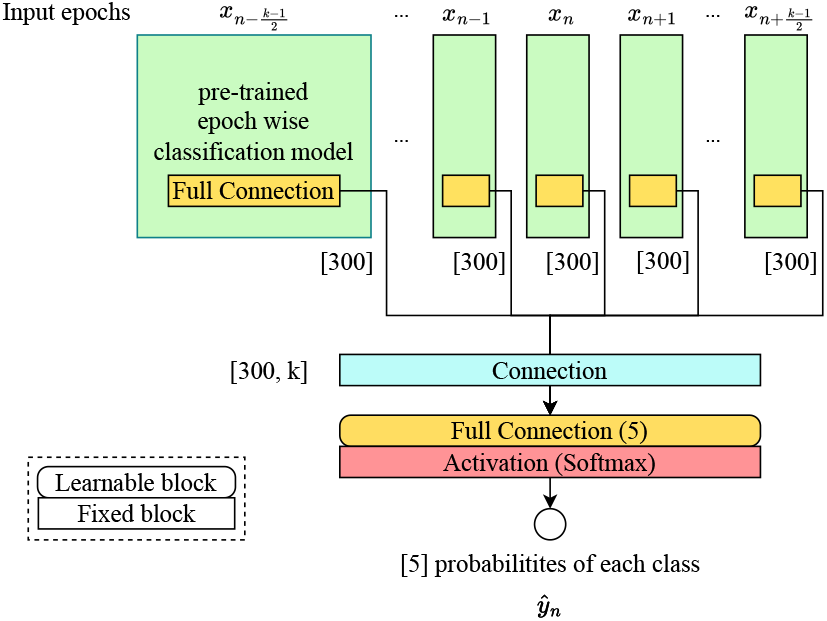
Transfer learning in sequential epoch data. The pretrained epoch-wise classification model is applied to *k* consecutive epochs, where *k* is an odd number ranging from 3 to 29. The outputs of the fully connected layers from the epoch-wise classification model are combined for the *k* epochs and passed through an additional fully connected layer with five units, followed by softmax activation to predict the probability of each sleep stage. The model is trained to predict the sleep stage of the central epoch in the sequence.

### C. LEARNING METHOD

Multiclass cross-entropy was used as the loss function. Learning terminated when the loss in the validation data had not improved over five consecutive iterations by using early stopping. Adam [29] was used as the optimizer, and the learning rate was set to 0.001, *β*_1_ = 0.9, *β*_2_ = 0.999, and *ϵ* = 10^*−*7^. The batch size was set to 32. For Sleep-EDF-20, leave-one-subject-out (20-fold) cross-validation was conducted to validate the model’s classification performance. For Sleep-EDF-78, all participants were randomly divided into 10 groups, and then subject-wise 10-fold cross-validation was conducted.

### D. EVALUATION METRICS

We used the following evaluation metrics of model performance: accuracy (ACC), macro-F1 score (MF1) [30], kappa coefficient (Cohen’s *kappa κ*) [31], and number of model parameters. Details of every metric are provided below.

ACC is the ratio of correctly predicted sleep stages to the total number of predictions. It provides an overall measure of the model performance but does not account for class imbalance. Sleep stage classification is a five-class classification problem, where every class is considered positive, and the other four classes are considered negative. The confusion matrix per class comprises four components: true positive (*TP*), false positive (*FP*), true negative (*TN*), and false negative (*FN*) rates.

MF1 is the unweighted mean of the F1 scores per class, with each F1 score being the harmonic mean of precision and recall. Precision is the ratio of true positive predictions to the total positive predictions, and recall is the ratio of true positive predictions to the total actual positives. MF1 treats each class equally regardless of its prevalence in a dataset, being suitable for imbalanced datasets. It emphasizes the performance of each class over the overall percentage of correct responses because it neglects the number of samples per class.

Cohen’s kappa measures the agreement between the predicted and actual sleep stages, considering the possibility of agreement occurring by chance. It is calculated as follows:

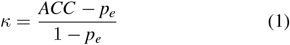

where *p*_*e*_ is the degree of coincidence given by

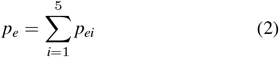

with the degree of coincidence per class being expressed as

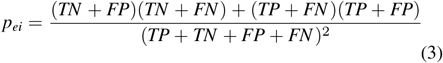

Kappa coefficients are interpreted according to their values, with a value of 0 indicating no agreement, and various ranges indicating slight (0.01–0.20), fair (0.21–0.40), moderate (0.41–0.60), substantial (0.61–0.80), and almost perfect (0.81–1.00) agreement [32]. We compared SleepSatelightFTC with existing models for sleep stage classification in terms of the abovementioned metrics and number of model parameters.

## III. RESULTS

### A. CLASSIFICATION PERFORMANCE ACCORDING TO NUMBER OF INPUT EPOCHS

ACC of the proposed SleepSatelightFTC model according to the number of input epochs for transfer learning is listed in Table 2. ACC on Sleep-EDF-20 was the highest at 85.73% for 25 input epochs, and ACC on Sleep-EDF-78 was the highest at 84.83% for 15 input epochs. We used the models with the highest ACC values to compare them with existing models.

**TABLE 2:** ACC according to number *k* of input epochs for transfer learning.

| # epochs<br>$k$ | Overall accuracy | |
| --- | --- | --- |
|  | Sleep-EDF-20 | Sleep-EDF-78 |
| 3 | 83.21 | 81.77 |
| 5 | 84.11 | 82.56 |
| 7 | 84.61 | 82.89 |
| 9 | 84.85 | 83.05 |
| 11 | 85.07 | 83.17 |
| 13 | 84.86 | 83.11 |
| 15 | 85.00 | **84.83** |
| 17 | 85.12 | 84.78 |
| 19 | 85.35 | 84.69 |
| 21 | 85.62 | 83.57 |
| 23 | 85.22 | 83.59 |
| 25 | **85.73** | 83.60 |
| 27 | 85.64 | 83.49 |
| 29 | 85.61 | 83.53 |

### B. CLASSIFICATION PERFORMANCE

A comparison of the classification performance between various models for sleep stage classification is presented in Table 3. The overall performances are given by ACC, MF1, and *κ*, and the class-wise performances are given by F1 scores. The performances for the existing models are those retrieved from the corresponding papers. The number of AttnSleep parameters is retrieved from [33]. SleepEEGNet, IITNet, DeepSleepNet-lite, CTCNet, WASR + LCNN, SleepTransformer, EEGSNet, and FFTCN were trained under the same conditions as SleepSatelightFTC. AttnSleep, TSA-Net, SeriesSleepNet, and TinySleepNet used weighted cross-entropy, and L-seqsleepnet used cross-entropy averaged over the sequence length. TinySleepNet used data augmentation during training.

**TABLE 3:**
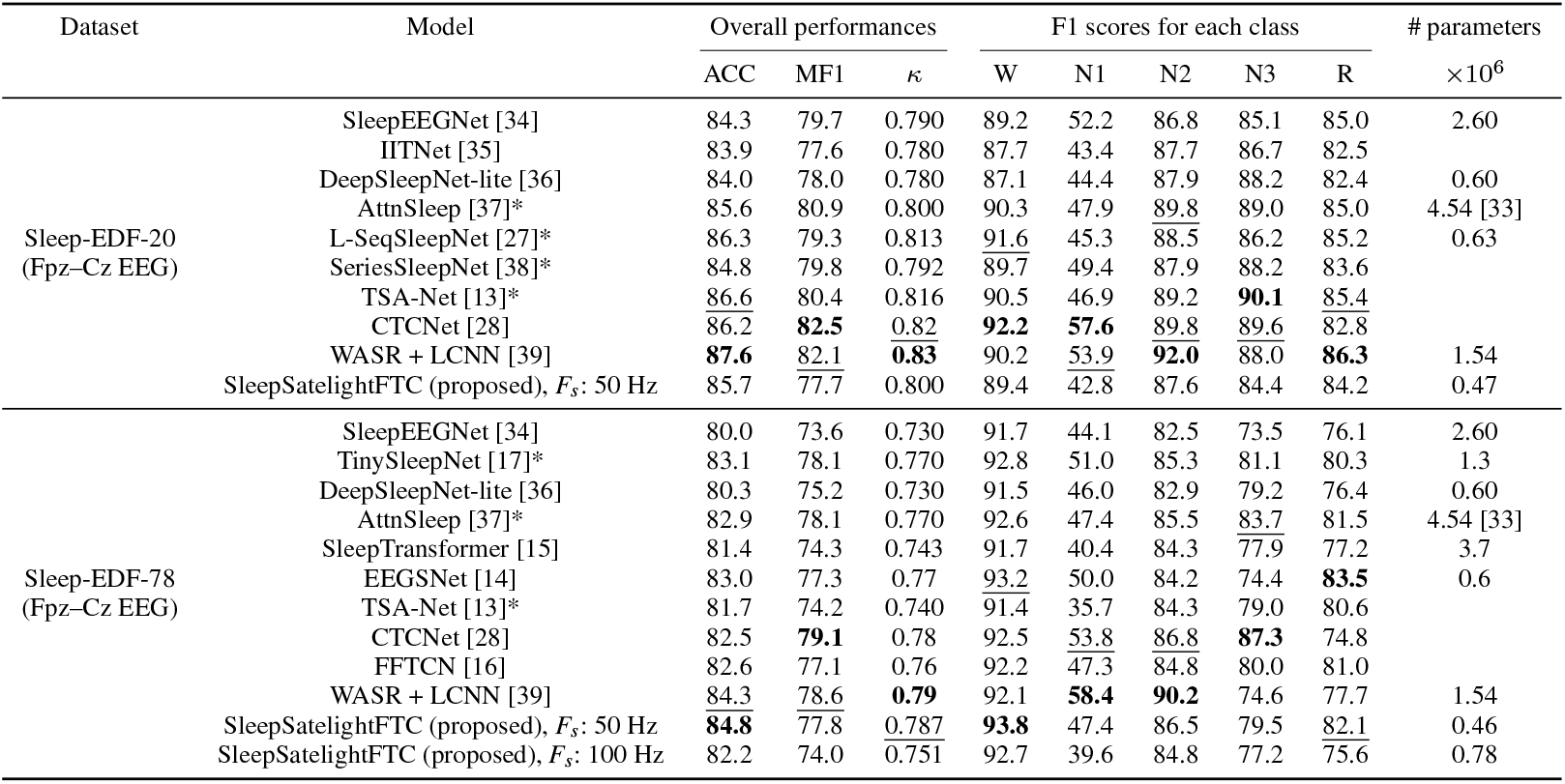
Sleep stage classification performance of evaluated models. *F*_*s*_ is the sampling frequency of input signals and * indicates methods that use different loss functions from that of SleepSatelightFTC. The number of AttnSleep parameters is retrieved from [33].

On the smaller Sleep-EDF-20, SleepSatelightFTC achieves an overall ACC of 85.7%, MF1 of 77.7%, and *κ* of 0.800. ACC and *κ* are nearly as high as the other models but slightly lower, with MF1 being particularly low.

On the larger Sleep-EDF-78, with EEG signals downsampled to a sampling frequency of 50 Hz, SleepSatelightFTC achieves an overall ACC of 84.8%, MF1 of 77.8%, and *κ* of 0.787. ACC and *κ* of our model are much higher than those of the state-of-the-art models. In addition, SleepSatelightFTC achieves the highest F1 scores in the classification of sleep stages W and N2. Additionally, when using the original EEG signals with a sampling frequency of 100 Hz, the model achieves an overall ACC of 82.2%, MF1 of 74.0%, and *κ* of 0.751.

### C. ABLATION STUDY

An ablation study confirmed that each component of the proposed SleepSatelightFTC model contributes to inference on Sleep-EDF-78. SleepSatelightFTC consists of time- and frequency-domain inputs as well as transfer learning. We evaluated the classification performance when one or two of these three components were removed from SleepSatelightFTC, obtaining the results listed in Table 4.

**TABLE 4:** Results of ablation study on Sleep-EDF-78.

| Model variant | Overall performances |  |  | F1 score of each class |  |  |  |  |
| --- | --- | --- | --- | --- | --- | --- | --- | --- |
| | ACC | MF1 | $\kappa$ | W | N1 | N2 | N3 | R |
| SleepSatelightFTC (time-domain input + frequency-domain input + transfer learning) | **84.8** | **77.8** | **0.787** | **93.8** | **47.4** | **86.5** | **79.5** | **82.1** |
| time-domain input + transfer learning | 82.5 | 74.4 | 0.753 | 92.6 | 40.8 | 85.1 | 77.1 | 76.2 |
| frequency-domain input + transfer learning | 81.0 | 72.8 | 0.732 | 90.7 | 38.5 | 84.1 | 76.6 | 74.2 |
| time-domain input + frequency-domain input | 78.9 | 69.7 | 0.704 | 90.5 | 30.0 | 83.5 | 78.5 | 66.1 |
| time-domain input | 78.3 | 67.8 | 0.693 | 90.5 | 25.6 | 82.9 | 76.5 | 63.4 |
| frequency-domain input | 75.7 | 64.6 | 0.654 | 86.1 | 20.5 | 81.7 | 74.8 | 60.2 |

When the frequency-domain input, time-domain input, and transfer learning are removed, ACC drops by 2.3%, 3.8%, and 5.9%, respectively. The F1 scores of sleep stage N3 are lower when the time- or frequency-domain inputs are removed than when transfer learning is removed. Furthermore, when the pair of frequency-domain input and transfer learning and the pair of time-domain input and transfer learning are removed, ACC drops by 6.5% and 9.1%, respectively. MF1 and *κ* also drop in these cases.

## IV. DISCUSSION

### A. COMPARISON OF PROPOSED AND EXISTING MODELS

The proposed SleepSatelightFTC model achieves higher ACC than existing models. In addition, the number of parameters in SleepSatelightFTC is 4.7 *×*10^5^, while that in the comparison models are 0.6–4.54 *×*10^6^, being approximately 1.3– 9.66 times larger than the number of parameters in SleepSatelightFTC. This can be attributed to the model architecture. Most existing models for sleep stage classification use raw EEG signals or spectrograms as inputs, extract epoch-wise features, and then consider contextual information before and after every evaluated epoch. In contrast, SleepSatelightFTC employs a parallel architecture that extracts features in both the time and frequency domains for subsequent integration. This approach allows the model to use information from multiple perspectives, effectively classifying sleep stages. Furthermore, applying self-attention to each domain enables the model to automatically learn and select essential features. The self-attention output shown in Figs. 4 and 5 confirms that the proposed model focuses on characteristic waveforms and frequency components, such as K-complexes and spindle waves. Additionally, introducing transfer learning using continuous epoch data enables judgments that consider the sleep context, thereby improving ACC in classification.

SleepSatelightFTC simplifies the context processing network of existing models by applying transfer learning to continuous epoch data. A single expert usually performs manual EEG-based sleep scoring. Therefore, existing models for sleep stage classification likely imitate the expert’s subjective evaluation [8]. By suppressing overlearning, lightweight models like SleepSatelightFTC are less likely to reflect subjective biases, possibly increasing the consistency and reliability of sleep stage classifications.

Given the lightweight architecture of SleepSatelightFTC, it is well-suited for real-time applications, particularly in clinical environments where quick decisions are necessary. The reduced number of parameters ensures faster inference times compared to larger models. In terms of computational efficiency, SleepSatelightFTC achieves a processing speed of 167 s per 1000 training steps, significantly outperforming other Transformer-based models such as SleepTransformer (308 s/1000 steps) [15] and L-SeqSleepNet (450 s/1000 steps) [27]. This substantial reduction in training time suggests that the model can also achieve faster inference speeds, making it a viable candidate for real-time or near-real-time deployment in sleep monitoring systems. While a more detailed speed analysis across different hardware platforms is necessary, these results indicate that SleepSatelightFTC provides a promising balance between computational efficiency and classification accuracy, making it highly suitable for practical applications.

SleepSatelight FTC classification performance on SleepEDF-20 was lower than in previous studies. The F1 scores per class for N1 and N3 were the lowest among the models we compared. This may be due to the limited number of samples for these stages in the smaller Sleep-EDF-20 dataset, which prevented the model from effectively learning their features. As a result, this contributed to the overall lower performance observed on this dataset.

SleepSatelightFTC achieves higher F1 scores for sleep stages N2 and W but lower F1 scores for sleep stage N1 than existing models on Sleep-EDF-20 and Sleep-EDF-78. This discrepancy may be due to the smaller number of epochs available for sleep stage N1 compared with those for sleep stages N2 and W on the Sleep-EDF Database Expanded. The use of weighted cross-entropy loss, as in AttnSleep, TSA-Net, and TinySleepNet, may improve the classification performance in sleep stages with scarce training data available.

The number of SleepSatelightFTC parameters increased by 1.8 when using EEG with a sampling frequency of 100 Hz, compared to a sampling frequency of 50 Hz. The overall performance decreased by 2.6% in the accuracy, 3.8% in the macro F1 score, and 0.036 in the Kappa coefficient. This is likely due to the fact that adding gamma waves as input, which are less relevant for determining sleep stage, prevented the model from learning the basis for inference. This result suggests that not including gamma waves as input may be effective in inferring sleep stages.

### B. CONTRIBUTION OF MODEL COMPONENTS TO INFERENCE

The proposed SleepSatelightFTC model achieves the highest ACC for 25 input epochs on Sleep-EDF-20 and 15 input epochs on Sleep-EDF-78, as listed in Table 2. In previous studies, context is considered from various epoch lengths, such as adjacent epochs [14], 15 epochs [40], and 100 epochs [41]. The ideal number of epochs to consider for the sleep context needs to be further analyzed, but the optimal number of epochs for the proposed model likely ranges from 15 to 25 epochs.

The ablation study results show that the performance declines the most when transfer learning is removed, followed by the removal of the time- and frequency-domain inputs. Hence, transfer learning as well as time- and frequency-domain inputs contribute to inference in that order. Each sleep stage typically lasts from a few to several tens of minutes, especially sleep stage N2, which accounts for approximately 45% of the total sleep time and is longer in late sleep stages [4]. During intervals of identical sleep stages, transfer learning is expected to compensate for out-of-context inferences.

### C. VISUALIZATION OF INFERENCE PROCESS

We visualized the inference process of SleepSatelightFTC using a dimensionality reduction method called uniform manifold approximation and projection [42]. The data distributions per class after time-domain input, frequency-domain input, self-attention layer output, epoch-wise inference, and transfer learning are shown in Fig. 3. Figs. 3a and 3b show that the model inputs are more coherently distributed by class in the frequency domain than in the time domain. On the other hand, Figs. 3b and 3c show that the self-attention layer outputs are more coherently distributed by class in the time domain than in the frequency domain. Overall, Fig. 3 shows the distribution becoming more clustered toward the latter half of the model. This suggests that self-attention may not necessarily be more effective for frequency features than for temporal features.

**FIGURE 3:**
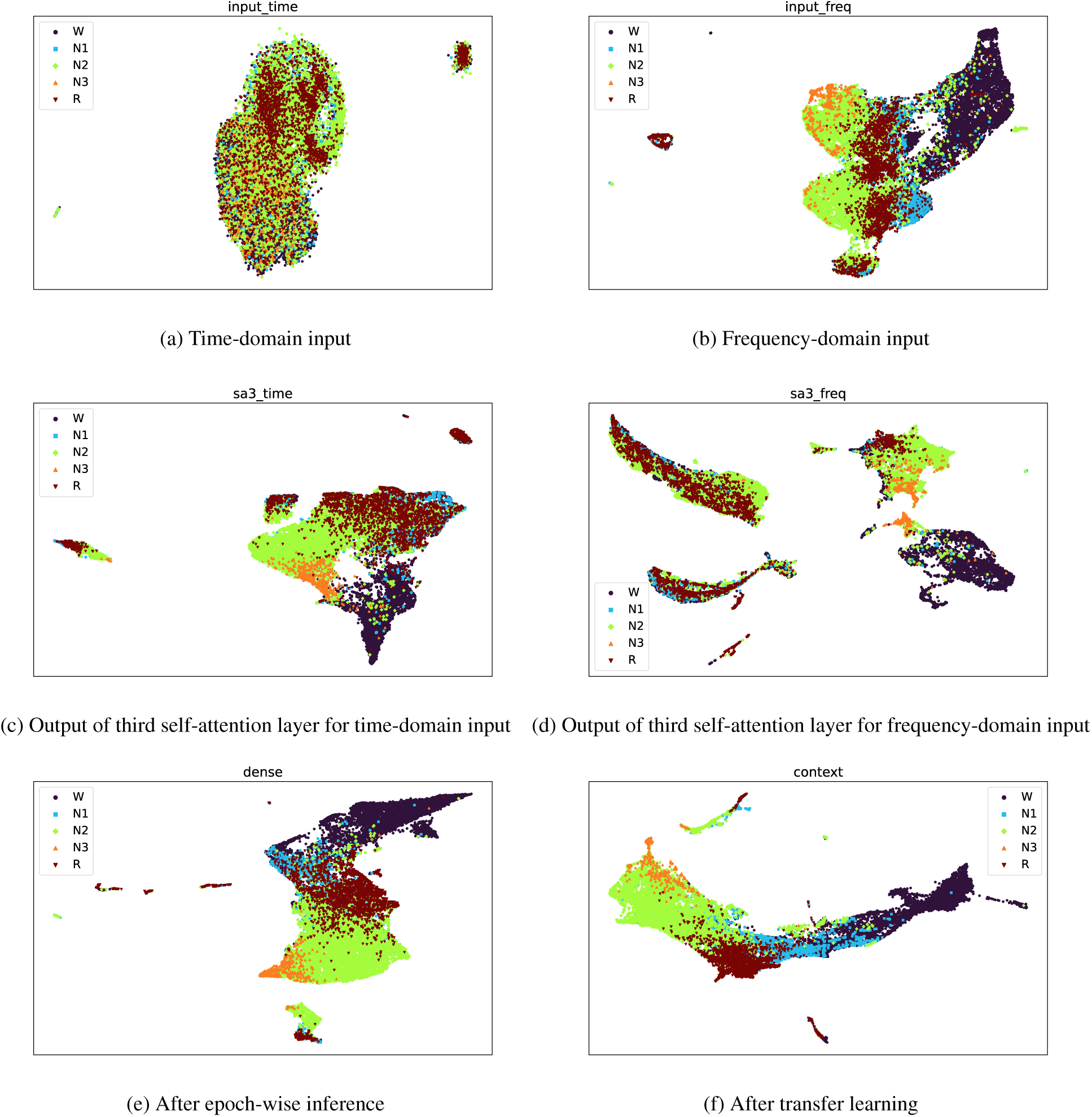
Visualization of inference process by uniform manifold approximation and projection.

### D. SELF-ATTENTION RESPONSES TO CHARACTERISTICS OF EACH SLEEP STAGE

We also created a heatmap of the output of the first self-attention layer for every input in SleepSatelightFTC. Because SleepSatelightFTC employs rectified linear unit activation, non-negative values of the output of the self-attention layer were set to 0. The output of the self-attention layer was averaged over time and normalized by using the function minmax_scale [43] from the preprocessing module of the scikit-learn library.

The model inputs and heatmaps of the self-attention layer outputs for an epoch correctly classified as sleep stage N2 are shown in Figs. 4 and 5. In the heatmap, self-attention responds strongly to a waveform that appears to be a K-complex, shown at the 15 s position in the EEG signal and the 12 Hz position in the amplitude spectrum.

**FIGURE 4:**
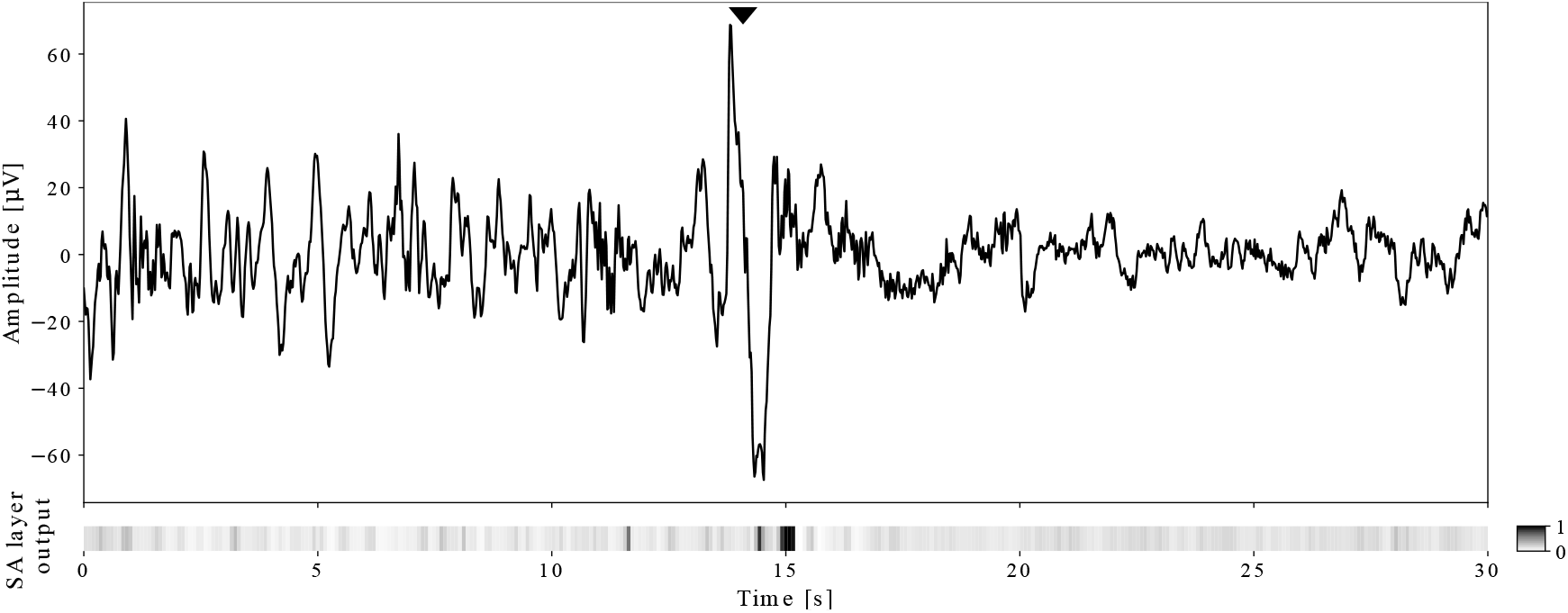
EEG signal for sleep stage N2 and its self-attention layer output heatmap. The heatmap shows the importance for classification of each timepoint in the EEG signal. The self-attention layer assigns higher importance to the waveform resembling a K-complex (▼), which is a characteristic feature of sleep stage N2.

**FIGURE 5:**
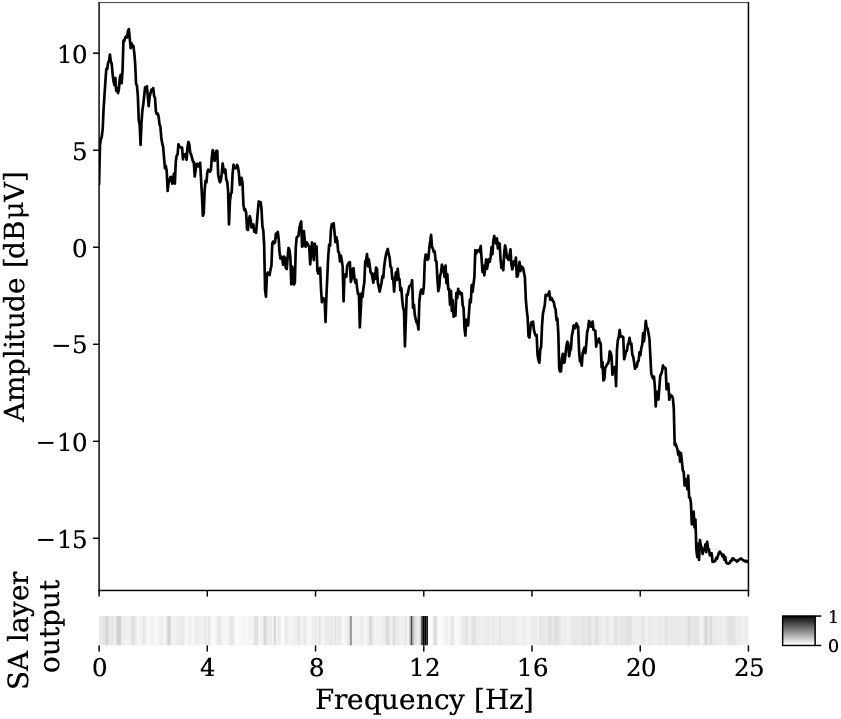
Amplitude spectrum of sleep stage N2 and its self-attention layer output heatmap. The heatmap shows the importance for classification o f e ach f requency component in the amplitude spectrum. The self-attention layer assigns higher importance to the 12 Hz frequency component, which is associated with sleep spindles, another characteristic feature of sleep stage N2.

The K-complex consists of a distinct negative sharp wave followed immediately by a positive component, observed especially in sleep stage N2 [3]. The K-complex duration is over 0.5 s, and the maximum amplitude is usually recorded in the frontal induction. In addition, spindle waves are features of sleep stages N2 and N3 characterized by a 12 Hz component in the frontal area [3]. The EEG signals of the Fpz–Cz channel considered in this study contain frontal EEG features. This suggests that SleepSatelightFTC learns K-complexes and spindle waves as features of sleep stage N2.

While recent Transformer-based models, such as Sleep-Transformer [15] and CTCNet [28], have demonstrated strong performance in sleep stage classification, SleepSatelightFTC offers several advantages in terms of model efficiency and interpretability. Transformer models generally rely on self-attention mechanisms applied across long temporal sequences, requiring substantial computational resources and a large number of parameters. In contrast, SleepSatelightFTC employs a lightweight self-attention architecture that focuses on epoch-wise time- and frequency-domain representations, maintaining interpretability while reducing computational costs. Specifically, CTCNet integrates a Transformer-based backbone to model sequential dependencies in sleep stages, but this comes at the cost of increased model complexity and computational requirements. Our results show that SleepSatelightFTC achieves comparable accuracy with significantly fewer parameters, making it a more practical choice for real-time applications. Additionally, the ability to visualize self-attention mechanisms in both time and frequency domains enhances its transparency, a critical aspect for clinical use.

While some Transformer-based models, such as MultiChannelSleepNet [11] or Cross-Modal Transformers [44], leverage multimodal data for improved classification performance, our study focuses on EEG-only models. Therefore, SleepSatelightFTC provides a more direct comparison to models like CTCNet and SleepTransformer, demonstrating the feasibility of an efficient, interpretable, and computationally lightweight sleep staging approach.

## V. CONCLUSION

We propose SleepSatelightFTC, a lightweight and interpretable deep learning model for EEG-based sleep stage classification that achieves higher ACC with fewer parameters than state-of-the-art models. The model employed self-attention to time- and frequency-domain inputs, raw EEG signals and amplitude spectrum, and transfer learning to sequential epochs to consider temporal context. The model interpretability through self-attention heatmaps, which highlight essential waveform features consistent with sleep scoring manuals, enhances the model accountability and allows experts to understand the reasons underlying its decisions and judge the validity of inference.

Nevertheless, our study has various limitations, such as using polysomnography data from only healthy subjects and not accounting for inter-rater variability in manual scoring. EEG patterns in individuals with sleep disorders may differ significantly from those of healthy subjects, affecting the model’s generalizability. In future work, we will evaluate the model performance based on data from patients with sleep disorders and explore domain adaptation techniques to address this issue. One potential approach is to fine-tune only the final fully connected layer of the trained model. Since sleep architecture varies between healthy individuals and those with sleep disorders, adapting the final classification layer while retaining the learned feature representations from healthy subjects may help mitigate domain discrepancies and improve model performance in clinical applications. Additionally, we plan to investigate methods to handle inter-rater variability and improve performance on underrepresented sleep stages.

Despite these limitations, SleepSatelightFTC demonstrates the potential of interpretable deep learning models for automatic sleep stage classification, which may substantially reduce the burden on sleep experts and improve the efficiency of sleep disorder diagnosis and treatment after further development and validation on diverse datasets.

## ACKNOWLEDGEMENTS

We thank Dr. Kosuke Fukumori of RIKEN AIP for his inspiration and advice. We would like to thank Editage (www.editage.jp) for English language editing.

